# Simulations reveal increased fluctuations in estrogen receptor-alpha conformation upon antagonist binding

**DOI:** 10.1101/061069

**Authors:** Ho Leung Ng

## Abstract

Molecular dynamics (MD) simulations have been used to model dynamic fluctuations in the structure of estrogen receptor-alpha (ER-α) upon binding to the natural agonist 17β-estradiol (E2) and to the active metabolite of the breast cancer drug and antagonist, 4-hydroxytamoxifen (OHT). We present the most extensive MD simulations to date of ER-α with over 1 μs of combined simulations for the monomer and dimer forms. Simulations reveal that the antagonist-bound complex includes significant fluctuations while the agonist-bound complex is tightly restrained. OHT increases dynamic disorder in the loops located to either side of the tail H12 helix; H12 has been associated with the activation status of ER-α. We also report that fluctuations near H12 lead to greater conformational variation in the binding mode of the ethylamine tail of OHT. Both the agonist and antagonist conformations are stable throughout the 240 ns simulations, supporting the hypothesis that there are no transitions between these two states or into intermediate states. The stable position of H12 in the OHT-bound conformation suggests that OHT stabilizes a well-defined antagonist conformational ensemble rather than merely blocking the agonist-driven activation of ER-α. Simultaneously, the increased dynamic properties of the OHT-bound complex is a potential source of binding entropy.

## Introduction

Estrogen receptor alpha (ER-α) is a transcription factor that mediates the primary physiological response to estrogens. It is a member of the nuclear hormone receptor family, which includes receptors for hormones such as thyroid hormone, androgens, and glucocorticoids [1,2]. Nuclear hormone receptors play a central role in mediating and regulating cell growth and death, development, metabolism, and immune responses. They are the targets of drugs for treating cancer, diabetes, inflammation, and autoimmune diseases [3,4]. Clinically used ER-α agonists include estrogen derivatives for contraception and menopausal symptoms. Drugs with mixed tissue-dependent antagonist and agonist effects are known as selective estrogen receptor modulators (SERMs) and include tamoxifen (used to treat breast cancer), clomiphene (used to treat infertility), and raloxifene (used to treat osteoporosis) [5,6].

ER-α, like other nuclear hormone receptors, contains three structural domains, an N-terminal domain that mediates dimerization and interacts with co-regulator proteins to promote gene expression, a DNA binding domain, and a C-terminal ligand binding domain (LBD) [7]. The crystal structure of full-length ER-α has not yet been determined; however, recent breakthroughs provided crystal structures of intact nuclear hormone receptor complexes bound to DNA that reveal close allosteric interactions between the domains [8].

Ligand interactions are entirely confined to the LBDs. LBDs have been the primary focus of structural studies and drug discovery efforts [9]. Over a hundred crystal structures of LBDs from nuclear hormone receptors bound to ligands have been published including dozens with the ER-α LBD co-crystallized with both agonists and SERMs. The active and inhibited states are associated with two different conformations of the C-terminal H12 helix (residues 538-548) through a “mouse trap” mechanism (Fig. 1) [2]. In the structure of the LBD bound to estradiol (E2), an agonist, H12 closes over the ligand to form part of the interaction surface with coactivators. When bound to antagonist (as in the structure of the LBD bound to SERM 4-hydroxytamoxifen (OHT)), H12 extends away from the ligand and occupies the co-activator binding site. Antagonists are generally larger than agonists (Fig. 2), and their bulk prevents H12 from adopting the active conformation.

**Fig. 1.**
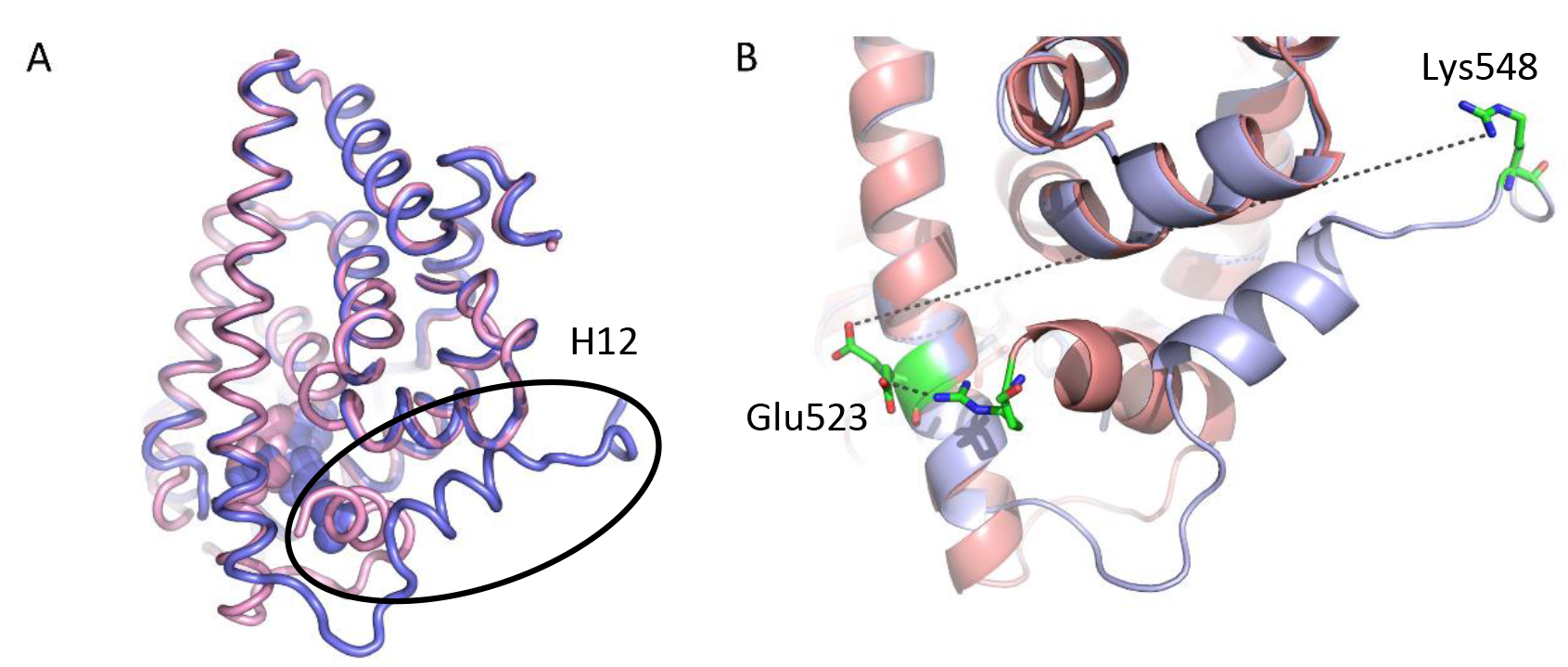
Crystal structures of ER-α bound to agonist and antagonist. A) Helix H12 (circled) adopts a closed conformation in the agonist (E2) bound structure (pink, PDB 1ere, 3.1 Å). Helix H12 extends away from the ligand in the antagonist (OHT) bound structure (blue, PDB 3ert, 1.9 Å). B) The agonist and antagonist bound conformations are distinguished by the distance between Glu523 and Lys548, which is short in the agonist-bound conformation (pink) and long in the antagonist-bound structure (blue).

**Fig. 2.**
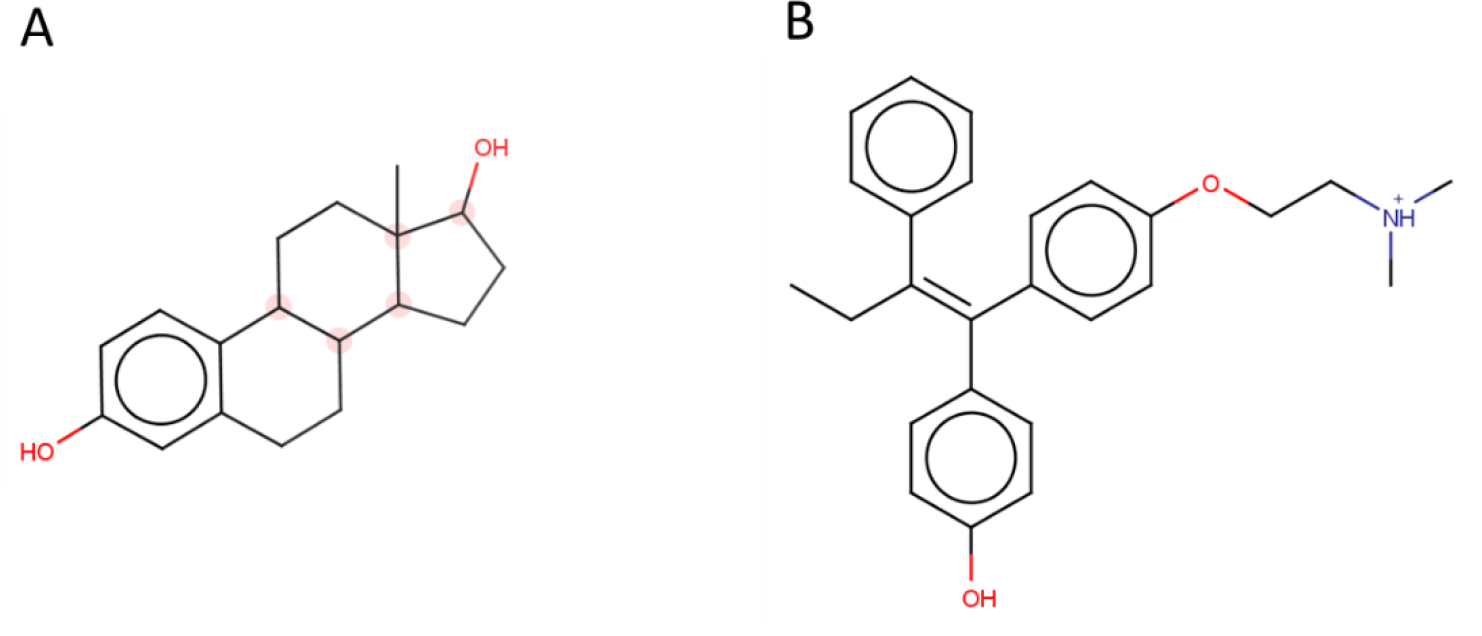
Structures of ER-α ligands. A) Physiological agonist 17-β-estradiol (E2). B) SERM antagonist 4-hydroxytamoxifen (OHT).

Recently, a number of studies have described surprising plasticity in the ER-α ligand binding site, supporting a link between distinct binding orientations and intermediate output states between full agonist and antagonist activities [10,11]. Molecular dynamics (MD) simulations have yielded further details into the dynamic properties of the agonist- and antagonist-bound conformations [12–17]. These studies indicate that the OHT-bound conformation is compatible with the docking of 11 other antagonists [14], suggest potential mechanisms for ligand release [15,16,18], demonstrate that the agonist-bound conformation is locked by co-activator peptide [12], and suggest the trajectory of the transition between the apo and agonist-bound conformations [12].

We hypothesized that an antagonist-bound complex would show greater dynamic activity than the agonist-bound complex. An agonist must lock the receptor into an active state to bind the co-activator, whereas an antagonist merely needs to interfere with the process. Here we present the results of the most extensive molecular dynamics calculations to date on ER-α with both agonist (E2) and antagonist (OHT) ligands, in monomer and dimer forms. Each monomer system was simulated in three parallel, independent runs for 240 ns. The dimers were simulated in single runs for 240 ns. Our calculations support the hypothesis that the antagonist complex is more dynamic than the agonist-bound conformation and identify receptor regions with increased fluctuation.

## Methods

The X-ray crystal structures of human ER-α with the physiological agonist, 17β-estradiol, (PDB 1ere) [19] and with the synthetic antagonist, 4-hydroxytamoxifen, (PDB 3ert) [20] were used as initial models for energy minimization and MD calculations. PDB 3ert is a high-resolution crystal structure with diffraction data to 1.9 Å, whereas PDB 1ere is based on a crystal that diffracted to 3.1 Å. Residues missing from the ligand binding domain, such as flexible loops, were modelled with Modeller [21] and Chimera [22]. All solvent and ion atoms located in the crystal structures were removed from the models for further calculations. Ligands were parameterized for the General Amber Force Field (GAFF) [23] using LEaP and Antechamber [24] from AmberTools 15 [25]. Hydrogen atoms missing from the crystal structures were added by LEaP. Each receptor-ligand complex was solvated in a rectangular box of TIP3P water molecules [26] in LEaP extending 10 Å from the complex in 0.15 M NaCl.

The solvated complexes were minimized by PMEMD in Amber 14 using the ff14SB [27] and GAFF force fields with the particle mesh Ewald method [28] with an 8 Å cutoff. Minimization was performed over three cycles in which the atomic coordinates were harmonically restrained with a weight of 5.0, then a weight of 1.0, and finally, were unrestrained. Each cycle included 100 steps of minimization by steepest descent and 900 steps by conjugate gradient.

Energy minimized structures were then equilibrated in MD simulations using the CUDA version of Amber PMEMD to support acceleration with NVIDIA graphics processing units [29,30]. Equilibration was performed in three cycles of 50,000 steps each, with a timestep of 1 fs, at constant pressure using a Berendsen barostat [31]. Temperature was maintained at 298 K with Langevin dynamics. In the equilibration cycles, the atomic coordinates were harmonically restrained with a weight of 5.0, a weight of 1.0, and, finally, a weight of 0.1.

Following equilibration, MD production runs of 240 ns were performed with Amber PMEMD with CUDA acceleration with constant pressure and a temperature of 298 K. Atomic coordinates were not restrained. The SHAKE algorithm was used to enable timesteps of 2 fs [32]. Different random number seeds were used for independent trajectories.

MD trajectories were analyzed with VMD 1.9.2 [33] and the CPPTRAJ tools from AmberTools 15 [34]. Protein structure figures were drawn with PyMol (Schrödinger, LLC). Root mean square deviations (RMSDs) of protein structures were calculated against the initial structure after the equilibration phase of molecular dynamics simulations using CPPTRAJ. Trajectories are available for download at holeungng.com.

## Results and discussion

We performed three independent MD simulations for monomer ER-α bound to E2 and bound to OHT using Amber14 accelerated with Nvidia GPUs, with each run lasting 240 ns [25,29,30].Simulations started from crystal structures of ER-α with these ligands. These are the longest MD simulations reported to date with ER-α. We performed the simulations with one of the most current and accurate force fields, ff14SB [27], with explicit solvent. This is also the first report of parallel simulations with ER-α, which allowed for more thorough configurational sampling and assessment of the significance and reproducibility of results. Our longer simulations showed that the calculated RMSD for Ca atoms in ER-α monomers did not reach a stable value until 20 ns (Fig. 3), longer than the total simulation time, 5 ns, of the most thorough MD study of ER-α published previously [12]. The mass-weighted radii of gyration of both E2 and OHT complexes converge at 18.7 and 18.9 Å during the simulations (Supplementary Fig. 1). The gyration radii of the E2 complex is stable from the beginning but that of the OHT complex does not plateau until 60 ns into the simulation. The stability of the radii of gyration for all the simulations supports the structural stability of the complexes under the trajectory parameters. Throughout all six runs, both E2 and OHT remained in the binding site, as demonstrated by the distance between the hydroxyl oxygen OE2 in ER-α Glu353 and the closest oxygen atom in the ligands (O3 for E2 and O4 for OHT) (Supplementary Fig. 2). In each of the three MD runs with OHT, the Ca RMSDs (average and maximum RMSD = 2.6 and 4.0 Å) were larger than those in runs with E2 (average and maximum RMSD = 2.1 and 3.3 Å). The increased fluctuations of the OHT complex relative to the E2 complex was also observed in the shorter 5-ns MD calculations reported by Celik et al., which showed a maximum RMSD of 3.5 Å for OHT vs 2.0 Å for E2 [12]. Our data support that longer trajectories are important when evaluating the significance of fluctuations as large RMSD fluctuations of up to 1.5 Å occurred within 10-ns time frames in both the E2 and OHT trajectories.

**Fig. 3.**
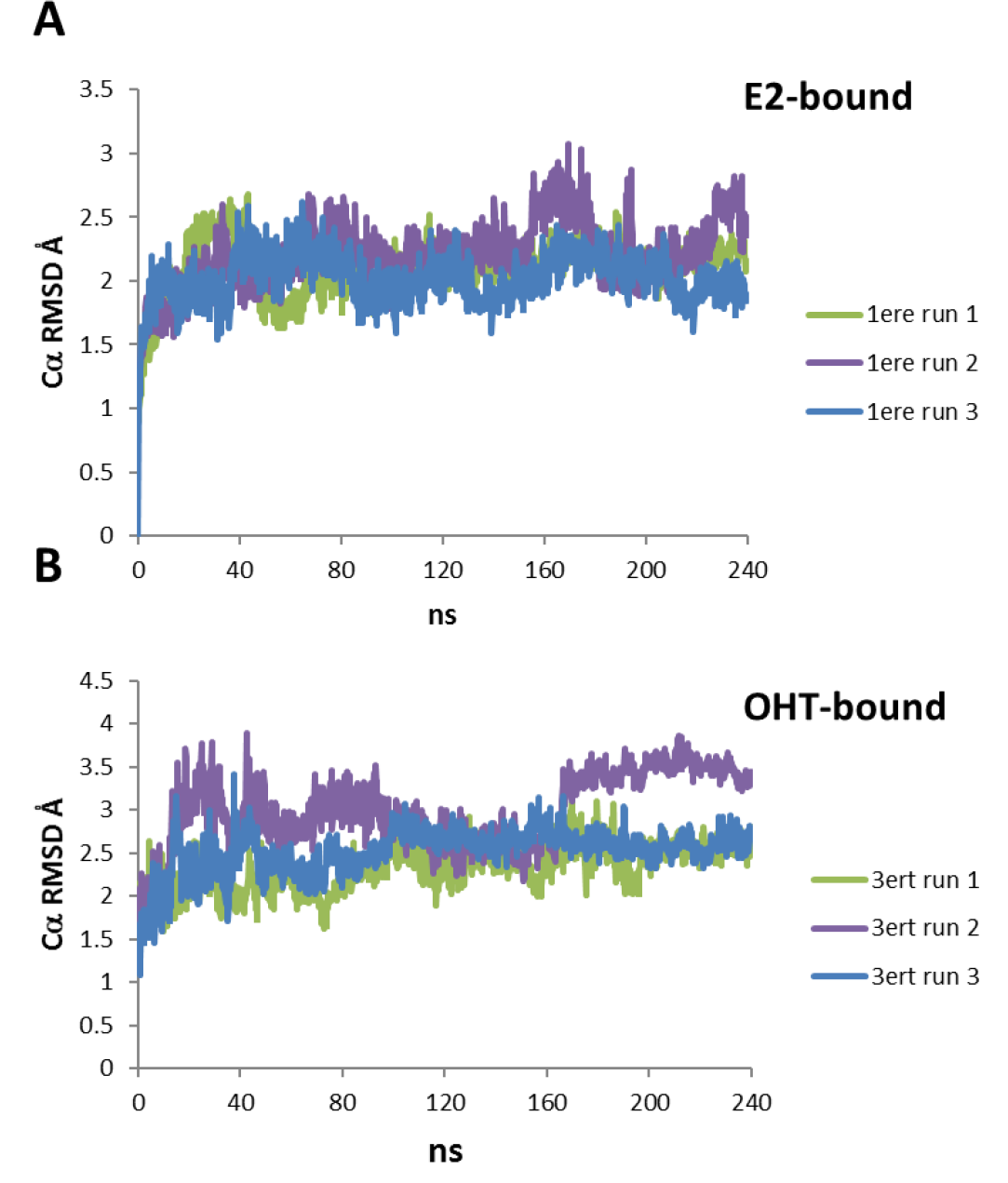
RMSD for Cα atoms throughout the MD simulations of ER-α with (A) E2 and (B) OHT.

Previous molecular dynamics studies focused on the monomeric form of ER-α because it was expected that binding and dimerization were independent events due to the remote distance of helix H12 from the dimerization interface [12]. However, data was not presented to support this assumption. We have performed 240 ns of molecular dynamics simulations of ER-α dimers with E2 and OHT to test this hypothesis. The Ca RMSDs for ER-α dimer with E2 plateaued at 2.1 Å after 60 ns, behaving similarly to the monomer (Supplementary Fig. 3). In contrast, the Ca RMSDs for ER-α dimer with OHT did not plateau but gradually increased over the simulation. At 240 ns, the Ca RMSD for ER-α dimer with OHT was 3.1 Å. Significantly longer simulations are probably needed to observe RMSD stability of the OHT dimer complex. Nevertheless, as with the monomers, the OHT dimer complex showed larger fluctuations than the E2 dimer. Both the E2 and OHT dimers show stable radii of gyration through the simulations (Supplementary Fig. 4), supporting the structural integrity of the dimers.

Analysis of root mean squared fluctuations (RMSFs) by residue in the monomers revealed that the increased dynamics of the OHT-complex compared to that of the E2-complex results from movement located exclusively in two regions, residues 526-535 and 545-550, corresponding to the loops that contact the ends of the terminal helix H12 (Fig. 4). Fluctuations in the loops surrounding H12 were also observed in a MD simulation that started with ER-α in the OHT-bound conformation but not when OHT was removed prior to the MD run [13]. Surprisingly, fluctuations in helixes H11 (residues 497-530) and H12 were similar in magnitude between the E2- and OHT-bound receptor trajectories despite significant conformational differences. This may reflect the high affinity of ER-α for each of these ligands. We also observed high dynamic activity in a loop preceding H9 (residues 160-166) in one of the three runs of ER-α with E2 (Fig. 4A). The increased fluctuations in this region was also observed previously in a 5 ns simulation of ER-α with E2 [35] and a 10 ns simulation with the mixed agonist/antagonist resveratrol [36]. Simulations of the dimers show the same regions with increased fluctuations as the monomers, further supporting that these dynamic effects are due to ligand binding (Supplementary Fig. 5). As the dimer simulations do not appear to provide significant additional information to the monomer simulations, the rest of our analysis focuses only on the monomers.

Analyses of interatomic distances further support that transitions do not occur between the agonist- and antagonist-bound conformations during the simulations of the monomers. The two conformations are easily characterized by the distance between the Leu544 in H12 and adjacent residues in either the agonist- and antagonist-bound states. The position of H12 was tracked in the E2-bound conformation by the distance between the centers of mass of Tyr526 and Leu544. In the crystal structure; this distance is 6.9 Å. Over the three trajectories, the mean distance between these two atoms was 7.0 Å (Fig. 5A). The position of H12 was tracked in the OHT-bound conformation by the distance between the centers of mass of Ile358 and Leu544. In the crystal structure; this distance is 6.3 Å. Over the three trajectories, the mean distance between Ile358 and Leu544 was 6.3 Å (Fig. 5B). The stabilities of the agonist and antagonist conformations throughout our simulations support our hypothesis that there are no transitions between these two states and no well-defined intermediate states within 240 ns.

Snapshots of coordinates taken at 180 and 240 ns in the three MD runs with E2- and OHT-bound receptor monomers also support that the receptor-ligand complexes are stable over the simulated time scale (Fig. 6, Supplementary Fig. 6). There was little deviation in the coordinates of the E2-bound complex (Fig. 6A). Most residues of the OHT-bound complex also showed little deviation with the exception of loop regions, particularly those surrounding H12 as described previously (Fig. 6B). The C-terminal amino acids partially unravel only in the OHT-bound complex.

**Fig. 4.**
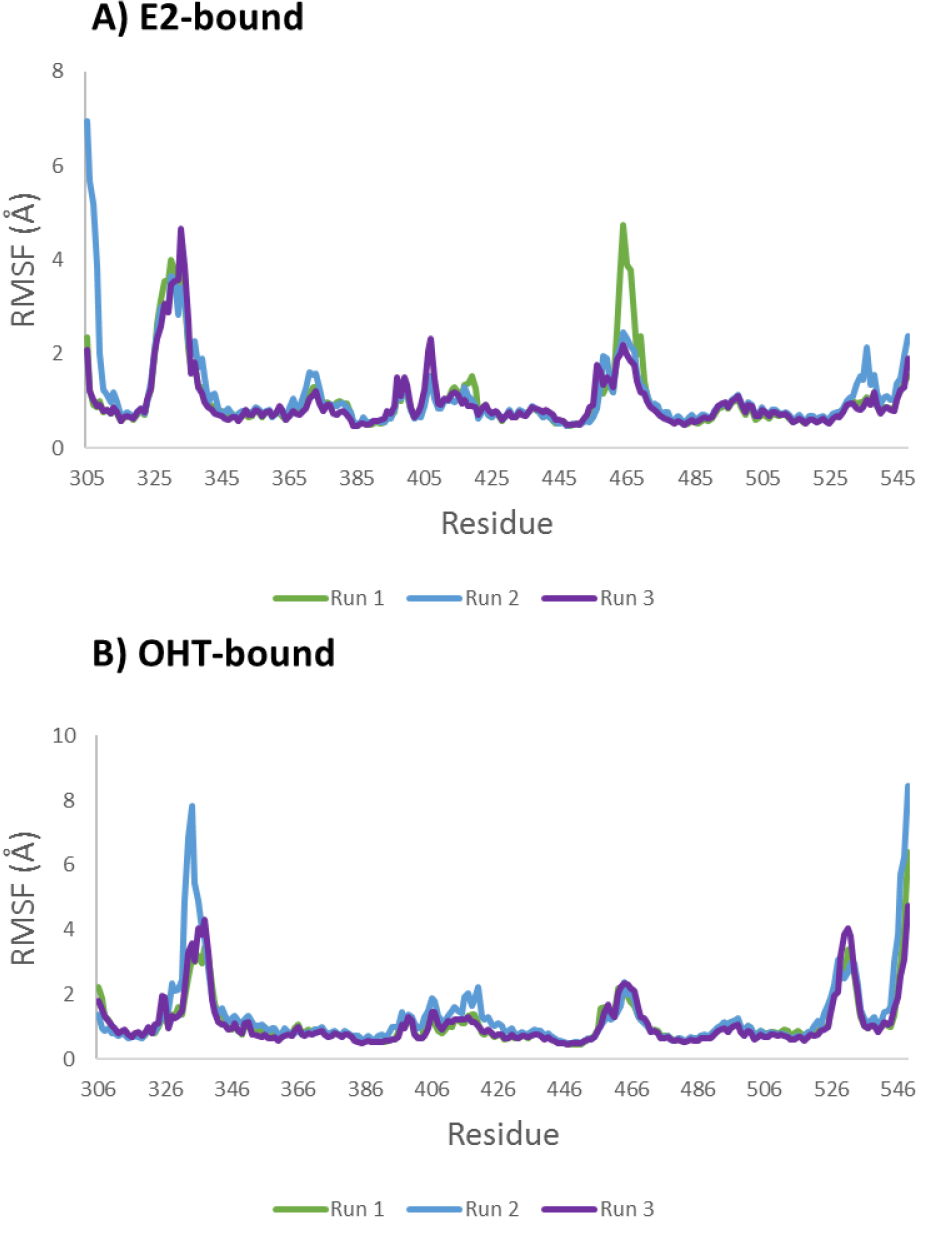
RMSF per residue for Cα atoms throughout the MD simulations of ER-α with (A) E2 and (B) OHT.

**Fig. 5.**
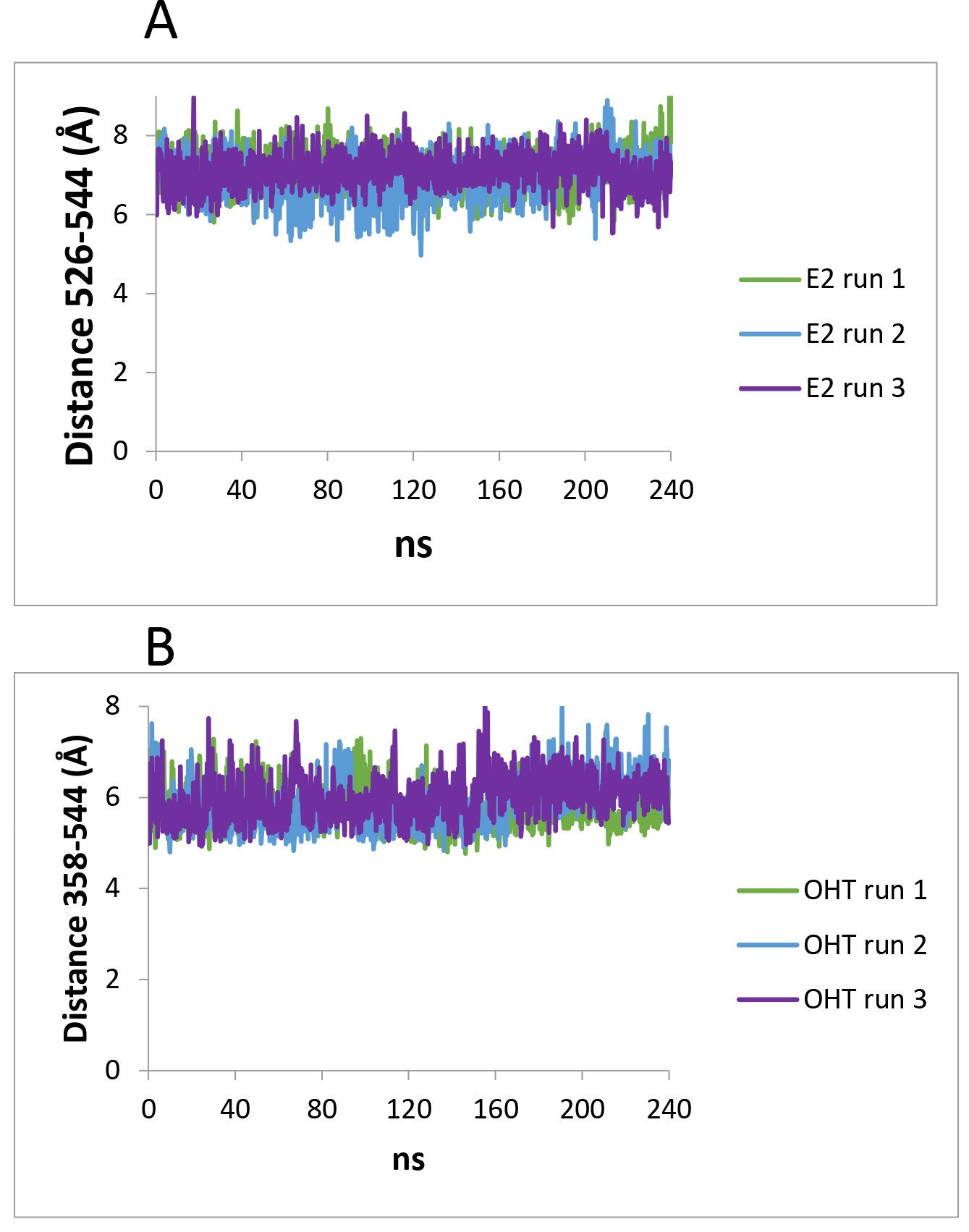
Agonist- and antagonist-bound conformations of H12. (A) H12 position in the E2-bound receptor defined by the distance between Tyr526 and Leu544. (B) H12 position in the OHT-bound receptor defined by the distance between Ile358 and Leu544.

**Fig. 6.**
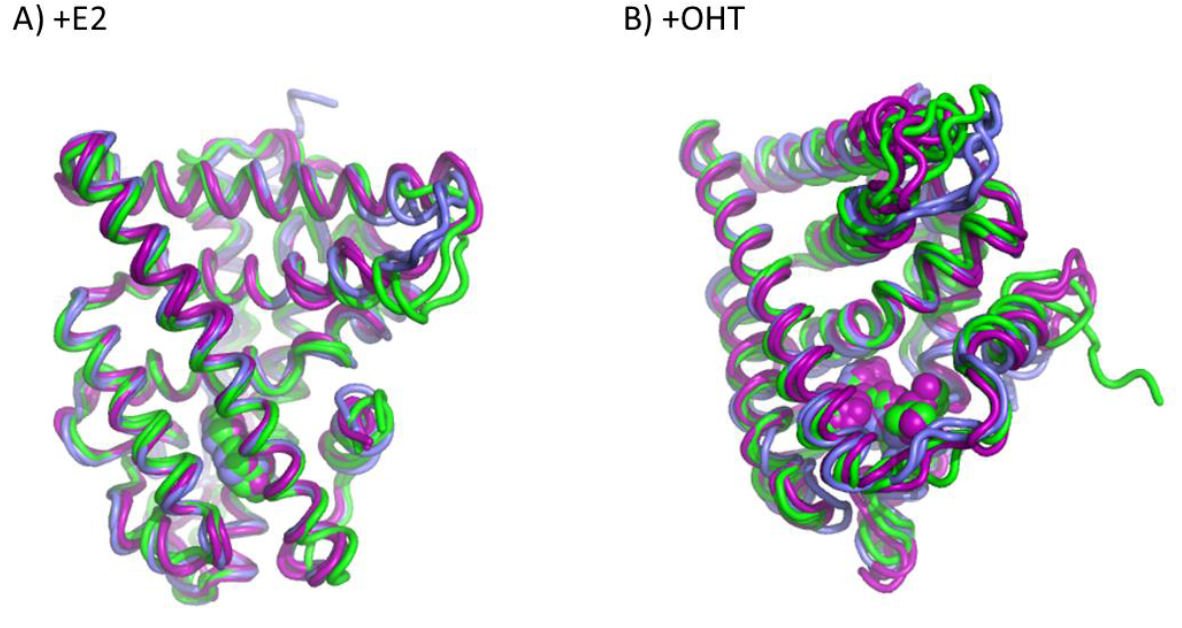
Coordinate snapshots taken at 180 and 240 ns from the three MD runs with ER-α bound to (A) E2 and (B) OHT. Green is from MD run 1, blue is from MD run 2, purple is from MD run 3

Increased conformational variation of the receptor translates to the ligand binding site. Coordinate snapshots taken at 180 and 240 ns from the three MD runs of monomers of both complexes were aligned based on Cα atoms. The conformation and binding orientation of E2 shows relatively little variation (Fig. 7A). In contrast, OHT shows more conformational and binding variation, especially in the ethylamine tail (Fig. 7B). We believe this is the first time the flexibility of bound OHT has been reported. This flexibility may be a source of entropy driving binding. OHT analogs with less conformational entropy in the bound state may be more potent antagonists. Alternatively, OHT may be bound more rigidly in a cellular environment in the presence of ER-α interacting proteins.

**Fig. 7.**
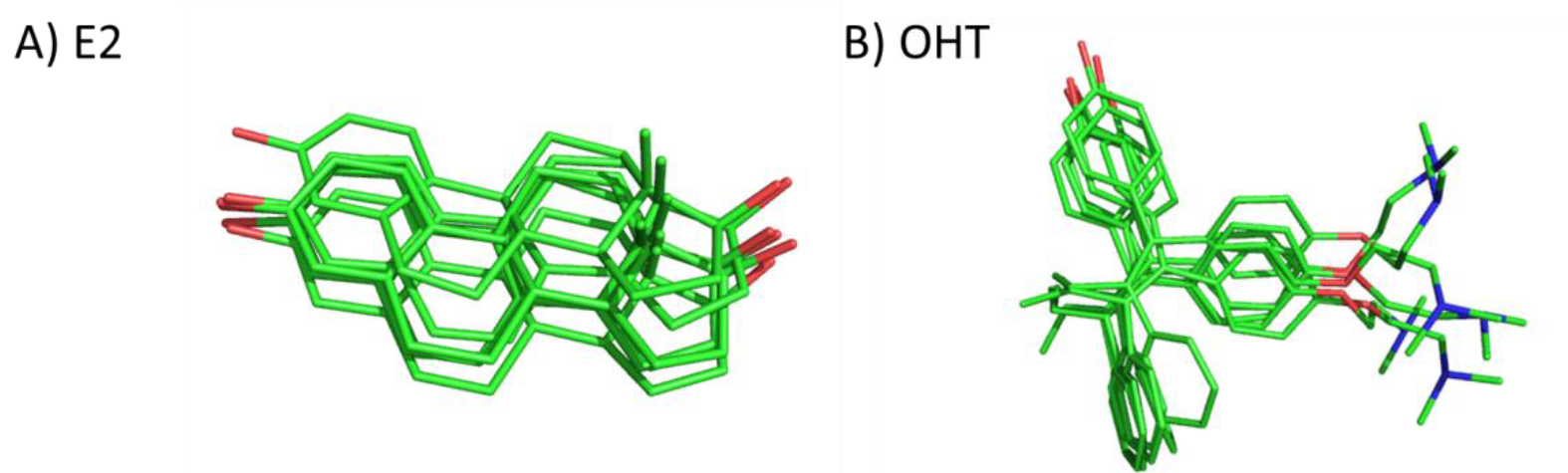
Ligand dynamics in the binding pocket for A) E2 and B) OHT. Atoms are colored green (carbon), red (oxygen), blue (nitrogen).

## Conclusions

We performed and analyzed the longest (240 ns) molecular dynamics simulations to date of ER-α with the agonist E2 and antagonist/SERM OHT. Our simulations indicate that simulations of at least 20 ns are required in order to ensure that the receptor-ligand complexes reach equilibration. Both the E2- and OHT-bound receptors were surprisingly stable throughout the course of multiple simulations. ER-α bound to OHT had much more significant structural fluctuations than did the receptor bound to the agonist. The fluctuations were primarily in the loops surrounding the key terminal helix H12. The fluctuating regions were conserved in simulations of dimeric ER-α with E2 and OHT, supporting that the dynamic differences are ligand driven. We also report that fluctuations near H12 lead to greater conformational variation in the binding mode of the ethylamine tail of OHT. Nevertheless, the well-defined conformation of H12 in the OHT-bound conformation supports the conclusion of an earlier study that OHT stabilizes a distinct antagonist conformational state rather than merely blocking the agonist-driven activation of ER-α [20]. Synthetic co-repressor peptides have been identified that bind to the OHT-bound complex [37], however, no natural co-repressors have been identified. The conformational stability of H12 in our MD trajectories is consistent with hydrogen/deuterium exchange (HDX) studies that demonstrated equal levels of exchange protection of H12 in the E2- and OHT-bound complexes [38]. The loops surrounding H12 that we have identified as the most dynamic were not studied by HDX. Intriguingly, OHT showed a unique HDX protection signature different from the SERM raloxifene, suggesting that our MD results are reflective of differential fluctuations in solution.

An unresolved question in estrogen receptor pharmacology is the molecular mechanism for partial agonism and mixed SERM activity. Clearly, the classical H12 on/off model is inadequate and does not account for graded and mixed ligand activity. Structural fluctuations explored in molecular dynamics simulations but not captured in static crystal structures may reveal deeper insights into these mechanism and identify features exploitable for drug discovery [39].

## Acknowledgements

The author is funded by NSF CAREER award 1350555.

